# IL-4 shaping glutamatergic synapse like structures for more mature iPSC derived neuron phenotypes

**DOI:** 10.1101/2023.12.05.570057

**Authors:** David Zimmermann, Clemens L. Schöpf, Georg Kern, Theodora Kalpachidou, Maximilian Zeidler, Michaela Kress

## Abstract

The prototypical pleiotropic anti-inflammatory cytokine IL-4 not only acts on immune cells but also has important roles in the nervous system mediating antinociception and neuroregeneration. Therefore, we explored the expression of IL-4, its receptors IL4RA and IL13RA1 and downstream signaling components together with morphological and functional assays as well as transcriptomic profiling in human nociceptors differentiating from induced pluripotent stem cells (iNocs).

IL-4 induced de novo formation of synaptic boutons immunoreactive for vesicular glutamate transporter vGLUT1 in iNocs which express both, components of the IL-4 receptor complex (IL-4RA and IL-13RA1) and signaling machinery (Jak1/2, STAT6, PKC isoforms, translation factor EiF4E) during differentiation. Pharmacological inhibition of these translational or cellular signaling components reduced the synaptogenic effect. IL-4 induced distinct transcriptomic changes associated with biological process ontologies for “neuron projection development”, “axonogenesis” and “synapse”, “cellular process involved in reproduction in multicellular organism”, “regulation of membrane potential” and “calcium ion transmembrane transport”. This partially reflected injury-induced transcriptional changes of mouse nerve injury models which contribute to regenerative processes in peripheral nerve but possibly also to reconnecting primary afferent neurons to their projections in the spinal dorsal horn and indicates a critical role for IL-4 in the control of developing and established neuronal networks.

## Introduction

Interleukin-4 (IL-4) is a prototypical member of the pleiotropic anti-inflammatory type 2 cytokine family which is generated and secreted by a variety of immune cells such as microglia, eosinophils or T cells, and not only acts back on immune cells such as microglia (Gieseck, Wilson, & Wynn, 2018; Mishra et al., 2021; Wynn, 2015; Zhu, 2015) but also has important roles in the nervous system (for review see (Gadani, Cronk, Norris, & Kipnis, 2012; Jiang, Cowan, Moonah, & Petri, 2018; Karp & Murray, 2012; Mamuladze & Kipnis, 2023; Quarta, Berneman, & Ponsaerts, 2020). IL-4 has garnered interest as a mediator of regeneration across multiple tissues including the central and peripheral nervous system where it for example exerts beneficial effects on neuronal survival, stem cell activity and even neurogenesis (Kiyota et al., 2010). In addition, IL-4 improves neuronal outgrowth and repair through a fast neuron-specific signaling pathway involving regulation of presynaptic vesicular glutamate transporter vGLUT1 in the central nervous system (cns) (Hanuscheck et al., 2020; Vogelaar et al., 2018). Although type 2 cytokines have been associated with pathological itch (Oetjen et al., 2017), the majority of findings suggest a homeostatic role of IL-4 and its receptors in neurogenesis, synaptic function, and neuronal regeneration after injury (Butovsky et al., 2006; Hanuscheck et al., 2022; Pan et al., 2022; Salvador, de Lima, & Kipnis, 2021). IL-4 secreting cells are recruited to the injury site within the cns or the injured nerve, and the secreted IL-4 promotes regeneration and repair (Pan et al., 2022; Walsh et al., 2015).

Upon peripheral nerve injury, IL-4 promotes peripheral nerve regeneration in rodents (Liao et al., 2019), and depletion of IL-4 severely compromises regenerative processes in the peripheral nervous system and the cns (Walsh et al., 2015; Zhang et al., 2019). Decreased levels of IL-4 are also associated with pain disorders or enhanced pain in humans and mechanical hypersensitivity in mice (Alexander, Perreault, Reichenberger, & Schwartzman, 2007; Schlereth, Drummond, & Birklein, 2014; Uçeyler, Eberle, Rolke, Birklein, & Sommer, 2007; Üçeyler, Topuzoğlu, Schiesser, Hahnenkamp, & Sommer, 2011; Uçeyler et al., 2006). Mice with a global depletion of IL-4, develop increased neuronal responses to noxious mechanical and chemical stimuli (Lemmer et al., 2015), and overexpression of IL-4 in dorsal root ganglion (DRG) neurons reduces these signs of neuropathic pain (Hao, Mata, Glorioso, & Fink, 2006). Also, spinal connections are frequently disturbed upon peripheral nerve injury via immune processes with losses of VGLUT1+ synapses (Campos, Barbosa-Silva, & Ribeiro-Resende, 2021; Gong, Hagopian, Holmes, Luo, & Xu, 2019; Y. Liu et al., 2017; Malet et al., 2013; Rotterman et al., 2019; Rotterman et al., 2023; Schmidtko et al., 2008; Takemura, Kobayashi, Kato, Yamaguchi, & Hori, 2018), and IL-4 has pro-regenerative as well as modulatory effects on synaptic structures (Castro et al., 2019).

IL-4 acts on its target cells by binding to heterodimeric IL-4RA/IL13-RA1 receptors (for review see (Shi, Song, Traub, Luxenhofer, & Kornmann, 2021; Wills-Karp & Finkelman, 2008). Multiple signaling components are involved including the activation of Janus kinases, STAT6, members of various cytosolic kinase families such as PKC isoforms, and even translation factors as well as mTOR (De Falco et al., 2015; Kalkman & Feuerbach, 2017; Shi et al., 2021; Sholl-Franco, Marques, Ferreira, & de Araujo, 2002; Vogelaar et al., 2018; Wills-Karp & Finkelman, 2008). IL-4 exerts analgesic effects and acts on nociceptive neurons (Cunha, Poole, Lorenzetti, Veiga, & Ferreira, 1999; Hao et al., 2006; Karam, Merckbawi, El-Kouba, Bazzi, & Bodman-Smith, 2013; Vale et al., 2003). However, mechanisms for IL-4 mediated actions favoring antinociception and peripheral nerve neuroregeneration and their translatability to human pathologies are controversially discussed in particular since type 2 cytokines and their receptors have been related to chronic itch and activate native and iPSC derived sensory neurons (Campion et al., 2019; Guo et al., 2022; Meng et al., 2021; Oetjen et al., 2017; Umehara et al., 2020). However, IL-4 may contribute beneficial analgesic and pro-regenerative effects and rescue injury induced vGLUT1 synapse loss of primary afferent neurons to inhibitory interneurons in the spinal cord.

Therefore, we set out to explore the expression of IL-4, its receptors IL-4RA and IL13RA1 and downstream signaling components together with morphological assays and unbiased transcriptomic profiling in differentiating human induced pluripotent stem cell (iPSC) derived nociceptors (iNocs). Differentially expressed genes were compared with single cell RNA sequencing data from mouse peripheral nerve injury models and comparative bioinformatics analysis revealed shared gene sets involved in axonogenesis and synapse formation.

## Results

### IL4/IL13RA1 expression and glutamatergic synapse formation

By analyzing publicly available single cell RNAseq data we addressed the expression of IL-4 receptor (IL-4RA) and IL13 receptor (IL13RA1) within different neuronal subtypes in human (GSE168243) and mouse (GSE155622, GSE154659) naïve DRG neurons. Integration of the neuronal sequencing data revealed conserved neuronal subtypes between the different datasets and species (Fig. 1 A,B). While cell numbers and expression levels of Il4RA and IL13RA1 varied between the datasets (Fig. 1C,D), similar expression patterns and mRNA proportions were found suggesting a robust expression of Il4RA and IL13RA1 in human and mouse DRG neurons. In order to explore the presence and role of IL-4 and its receptors in iPSC derived neuron differentiation or maturation we first retrieved trajectories for relevant subunits of the IL4/IL13 receptor complex in silico using the NOCICEPTRA data set and app (Zeidler et al., 2021). While IL-4 mRNA was not at all detectable in iPSCs or developing iNocs (data not shown), IL4RA and IL13RA1 mRNA was highly expressed already at the pluripotency stage, with expression levels declining until D16 followed by increasing expression of both receptor subunits but IL13RA1 rising to considerably higher expression levels than IL4RA (Fig. 1 E,F). In order to explore the importance of IL-4 for potential regenerative and synaptic processes in developing iNocs we exposed iNocs at D16 to a physiological concentration of IL-4 for 3h and 24h and performed immune staining for the vesicular glutamate transporter type 1 (vGLUT1) (Fig. 1G). Synaptic boutons were thresholded in the vGLUT1 channel and quantified using an integrated morphometry plugin of Metamorph^®^ (IMA-mask, methods) in order to assess all vGLUT1 clusters independent of their cellular localization. Upon IL-4 treatment, a significantly increased number of vGLUT1 positive puncta, were detected (Fig.1 G). Upon prolonged IL-4 treatment (24h) the number of vGLUT positive boutons was profoundly increased indicating that individual iNocs formed significantly more vGLUT1+ clusters, whereas the iNoc number remained stable suggesting an active role of IL-4 in de novo glutamatergic synapse formation (Fig. 1 H,I). vGLUT1 immunoreactivity was co-localized in synaptic boutons with immunoreactivity for the neuronal filament marker TUJ1 (Fig. 1J). To quantify the effect of IL-4 on glutamatergic synapse formation, additional TUJ1/vGLUT1 immune staining was performed. Both, the total number of vGLUT1 clusters as well as TUJ1+/vGLUT1+double positive synaptic boutons increased upon prolonged IL-4 treatment (Fig. 1K) but no significant changes were observed with other cytokines (Suppl. Fig. 1) indicating that the regulation of synaptic glutamatergic boutons might be specific for IL-4. Interestingly, iNocs also expressed mRNAs for a subset of ion channels of the AMPA and NMDA receptor family as well as metabotropic receptors for glutamate which seemed to be functional in all iNocs as observed in microfluorimetric Ca^2+^ measurements which showed strong responses to glutamate suggesting the formation of functional synapses due to IL-4 mediated development of glutamatergic presynaptic structures in human iNocs (Suppl. Fig. 2).

**Figure 1.**
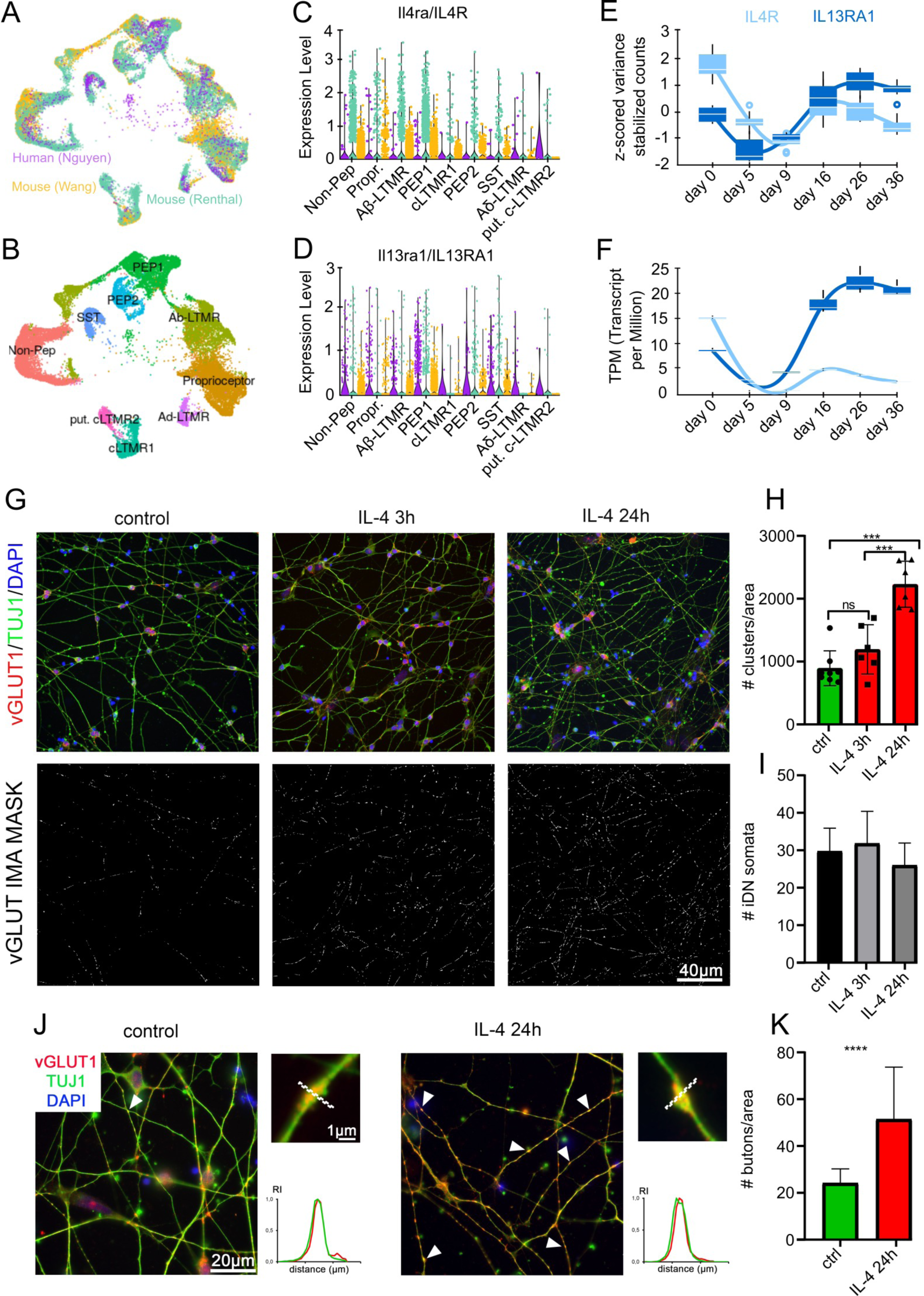
IL4RA/IL13RA1 expression in different nociceptive subtypes and IL-4 induced synapse formation in iDNs. (A) Human (GSE168243 - purple) and mouse (GSE154659 - yellow, GSE155622 - cyan) RNAseq datasets were integrated and (B) neuronal subpopulations were annotated according to marker genes Renthal et al. (C, D) Comparison of IL4RA and IL13RA1 mRNA expression along identified neuronal subtypes in human and mouse datasets: non-peptidergic neurons (Non-Pep), proprioceptors (Propr), Aβ low threshold mechanoreceptors (Aβ-LTMR), Tac1+/Gpx3+ peptidergic nociceptors (PEP1), Tac1+/Hpca+ peptidergic neurons (PEP2), Fam19a4+/Th+ C-fiber low threshold-mechanoreceptors (cLTMR), somatostatin positive pruriceptors (SST), Aδ low threshold mechanrecepotrs (Aδ-LTMR) and Fam19a4+/Th low putative cLTMR (put-cLTMR). (E, F) Both IL4RA and IL13RA1 were robustly expressed in developing iNocs as queried in NOCICEPTRA; normalized count trajectories and TPM expression levels were illustrated in DIV0-DIV36 iNocs. (G) iNocs predominantly developed into vGLUT1+ glutamatergic neurons and the number of glutamatergic synapses increased upon IL-4 treatment. IL-4 treatment was performed for 3h and 24h in iDNs. vGLUT1 positive boutons were detected in D16 iNocs and counterstained with TUJ1 (neurites) and DAPI (nuclei), magnifications highlight individual vGLUT1+ clusters on putative synaptic boutons, colocalization of vGLUT1 and TUJ1 fluorescent intensities were quantified by line scan analysis (H) Quantification of total vGLUT1^+^ clusters 24h after IL-4 treatment. (I) The number of iNoc somata did not change upon treatments. (J) Individual sprouting synaptic boutons were labeled by vGLUT1^+^/TUJ1^+^ co-staining. (K) Quantification of vGLUT1^+^/TUJ1^+^ glutamatergic boutons before and after 24h IL4 treatment. Upon IL-4 treatment the number of vGLUT positive boutons robustly increased, arrowheads, quantification (p<0.0001).

### Multiple pathways involved in IL-4 regulation of vGLUT/TUJ1 positive boutons

In order to assess potential signaling pathways involved in the IL-4 induced formation of glutamatergic boutons we queried the NOCICEPTRA database for signaling components downstream of IL-4R activation in silico and retrieved trajectories of Janus kinases JAK1 and JAK2, signal transducer and activator of transcription STAT6 and tyrosine kinase TYK2 expression which steadily increased after D09 whereas JAK3 and STAT3 expression levels remained low throughout the entire differentiation process (Fig. 2A). These findings suggest classical signaling pathways of IL-4 in iNocs upon exposure to the cytokine which itself was not expressed at relevant levels (data not shown). We next performed a pharmacological approach and co-applied small molecule inhibitors of key translational and transcriptional pathways components (Fig. 2A,B) to inhibit eukaryotic translation initiation factor eIF4E (EIF4Ei), STAT6 (STAT6i), mammalian target of rapamycin mTOR (RAP) or the protein kinase C PKC pathway (Bim1). Inhibitors were applied 1h before and together with IL-4 for 24h and the number of sprouting glutamatergic boutons was compared between IL-4 treated iNocs, vehicle treated controls and inhibitors applied together with IL-4. All of the inhibitors significantly (p<0.0001; Fig. 2C,D) reduced the number of glutamatergic boutons compared to IL-4 treated iNocs. Total numbers of cells were not affected by the treatments (Suppl. Fig. 3). Neurons treated with STAT6i showed even a lower number of vGLUT/TUJ1 positive boutons as compared to untreated control iNocs. These findings suggest that multiple classical signaling pathways appear to be involved in presynaptic synapse formation in iNocs, however also highlight a broader role of STAT6 since STAT6i not only affected IL-4 induced but also basic synapse formation.

**Figure 2.**
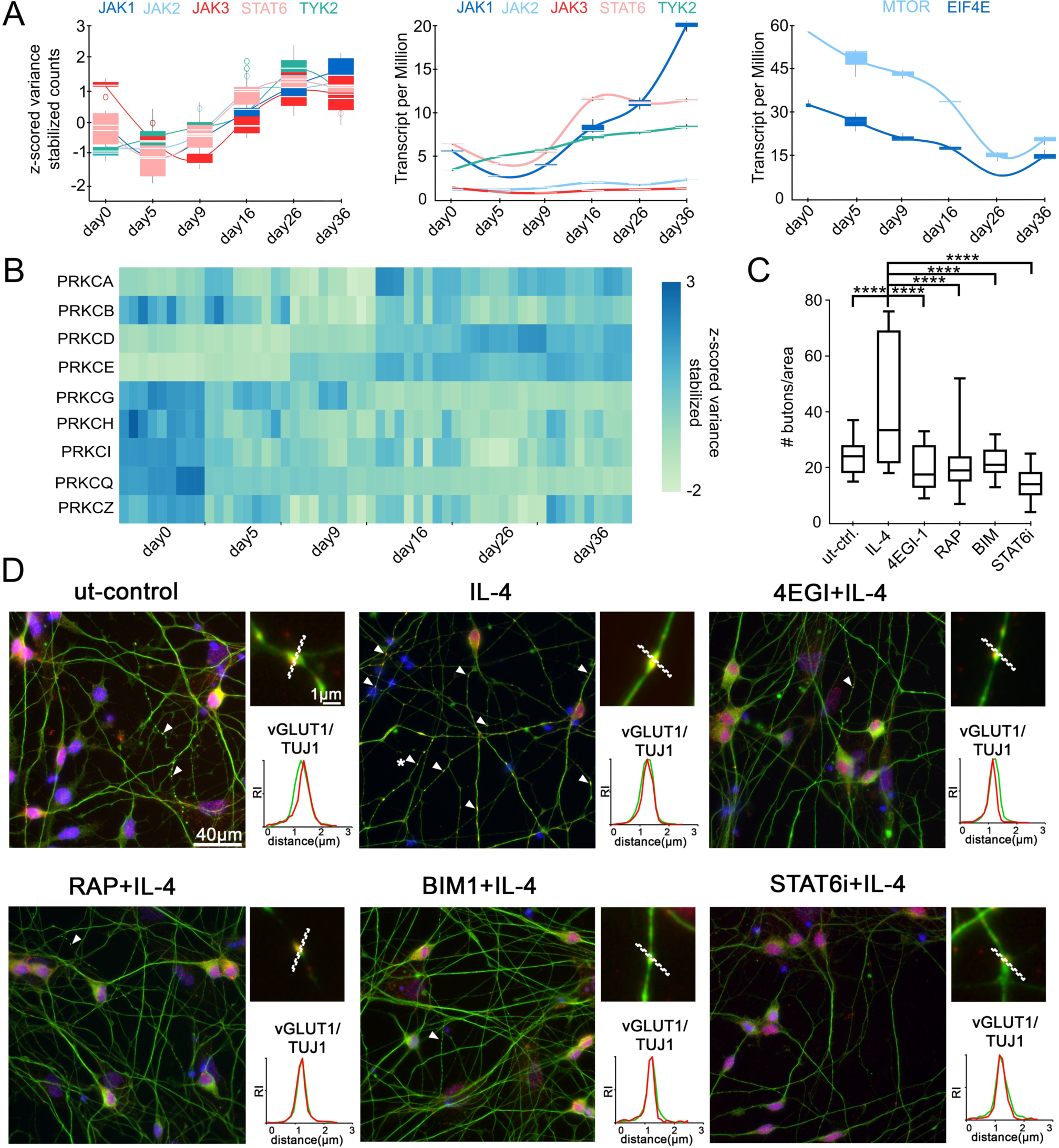
Multiple signaling pathways involved in IL-4 induced synapse formation in iNocs. (A) Expression of IL-4-downstream signaling components in developing iNocs: Normalized count trajectories and TPM expression levels of JAK1-3/STAT6/TYK2 as well as MTOR/eIF4E mRNA were assessed by the NOCICEPTRA app (B) Heatmap showing the expression of 9 different PKC isoforms during iDN development with prominent expression of PKCA-E at D16 during synaptic development; (C, D) Quantification of pathway inhibitors of synapse formation: vGLUT1^+^/TUJ1^+^ boutons per area were compared between ut-controls, 24h IL-4 (20ng) stimulated iDNs or 1h inhibitor pretreated iDNs following IL-4 stimulation (inhibitors: STAT6i (1µM), eIF4Ei (40µM), RAP (200nM), BISL (1µM)). IL-4 vs. ut control, p<0.0001, IL-4 vs. eIF4E+IL-4 (p<0.0001), IL-4 vs. RAP+IL-4 (p<0.0001), IL-4 vs. BIS+IL-4 (p<0.0001), IL-4 vs. STAT6i+IL-4 (p<0.0001).

### IL-4 regulated genes encoding for glutamatergic synapse organizing elements

In order to address transcriptomic changes induced by IL-4 treatment, we next performed RNA sequencing of 24h IL-4 treated and untreated iNocs (D16, Suppl. Fig. 4) which were separatable and groupable by their gene expression profiles after principal component analysis (PCA; Fig. 3A). Differential gene expression (DGE) analysis was performed, and the Benjamini-Hochberg algorithm was applied for false discovery rate (FDR) correction due to multiple testing. This analysis revealed 932 significantly (p_adj_<0.01) upregulated and 1577 significantly downregulated differentially expressed genes (DEGs) after IL-4 treatment (Fig. 3B). Gene ontology (GO) enrichment analysis for the significantly upregulated DEGs revealed the highest number of genes associated with biological process (BP) ontologies for neuron projection development (47 genes), axonogenesis (43 genes) and synapse organization (43 genes) (Fig. 3C). GO-analysis of the downregulated genes revealed the highest number of genes associated with BP-ontologies for cellular process involved in reproduction in multicellular organism (29 genes), regulation of membrane potential (29 genes) and calcium ion transmembrane transport (28 genes) (Fig. 3D). Analysis of 639 human genes linked to glutamatergic synapse ontology (cellular component GO:0098978) revealed 144 significantly (padj <0.05) upregulated and 46 downregulated genes (Supplementary Table S1) after IL-4 treatment. Of these genes, 17 upregulated and 5 downregulated genes can be exclusively linked to presynapse GO (GO:0098793), 42 upregulated and 15 downregulated genes were linked to both, pre-synapse and post-synapse GO (GO:0098794). Investigation of 89 human orthologs of GO regulation of synaptic transmission, glutamatergic (GO:051966) revealed 15 significantly up- and 10 significantly downregulated genes. In detail, *NRXN1*, *SERPINE2*, *NLGN1*, *CNR1*, *DGKI*, *NTRK2*, *STXBP1*, *GRIA4*, *SYT1*, *MYO5A*, *NGFR*, *ADCYAP1*, *CDH2*, *CACNG7*, and *GRIK1* were upregulated, and *GRM7*, *DRD3*, *RELN*, *ROR2*, *SHANK2*, *GRM5*, *GRIN2B*, *GRIN2A*, *CACNG3*, and *PLA2G6* were downregulated. The data indicate that IL-4 treatment in iNocs led to changes in transcript levels of proteins that are essential for the structure and function of synaptic glutamatergic connections and signaling.

**Figure 3.**
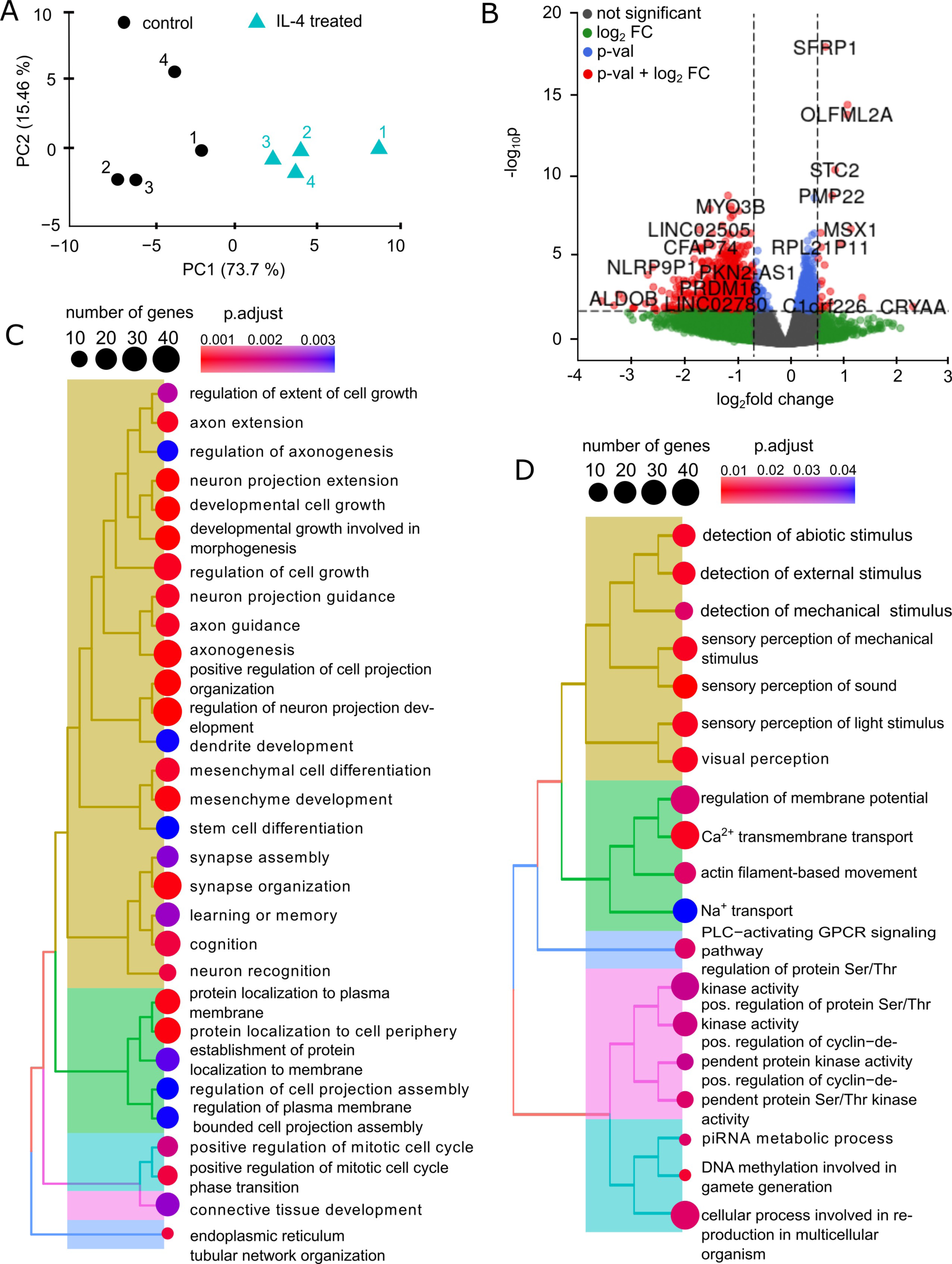
Transcriptional changes in iDNs after external IL-4 treatment. (A) RNA of four samples untreated iDNs and four samples 24h IL-4 (20ng)-treated IDNs was sequenced and principal component analysis (PCA) was applied to analyze the treatment effects showing a clear separation of treated (black circles) vs. untreated iNocs (cyan triangles) (B) Differential Gene Expression Analysis was performed between treated vs untreated samples. Differentially expressed genes after IL-4 treatment are represented in the volcano plot with threshold set to FDR corrected p-value ≤ 0.01 and |log_2_fold change| = 0.6. (C, D) Significant DEGs were ranked and gene ontology enrichment analysis was performed for (C) upregulated and (D) downregulated genes of biological process ontology.

### Deregulated genes overlapping with transcriptomic profiles of injured neurons

To address the biological importance of these findings in iNocs we exploited publicly available single cell RNAseq data sets from mouse dorsal root ganglion cells only since human injury data sets are currently unavailable (GSE155622, GSE154659). Recent findings in peripheral nerve injury models revealed injury specific cluster(s) of cells as early as 24h and up to 60 days after injury in mice (Renthal et al., 2020; Wang et al., 2021). We integrated single nuclei data before (naive) and after crush-injury (Crush), ScNT injury and single cell data before and after SNI (Fig. 4A) and focused on the transcriptional differences at 7 days after the injury since this time point was provided in all studies (Fig. 4B). A new cluster appeared in all data sets after injury, and the percentage of cells in this “injury” cluster increased from 2.5% of all sequenced cells at 12h to a peak percentage of 44.14% at 72h after injury (Supplementary Figure 4A). To investigate the potential role and the type of IL-4 signaling upon nerve injury, we analyzed the amount of Il4RA+/Il13RA1-, Il4RA-/Il13RA1+ and Il4RA+/Il13RA1+ expressing cells (Fig. 4C) as well as Il4ra and Il13ra1 expression levels (Supplementary Figure 6). Since the absolute number of cells was different between naïve and 7d post injury, we compared the relative number of cells and found an overall increase of Il4RA+/Il13RA1+ as well Il4RA-/Il13RA1+ characterized cells in all datasets and injury types. However, the relative amount of Il4RA+/Il13RA1-cells increased in the Crush (10.39 % before and 15.46 % after crush injury) and ScNT (10.39% before and 15.26 % after ScNT) data and decreased in the SNI data (21.59 % before and 17.36 % after SNI).

**Figure 4.**
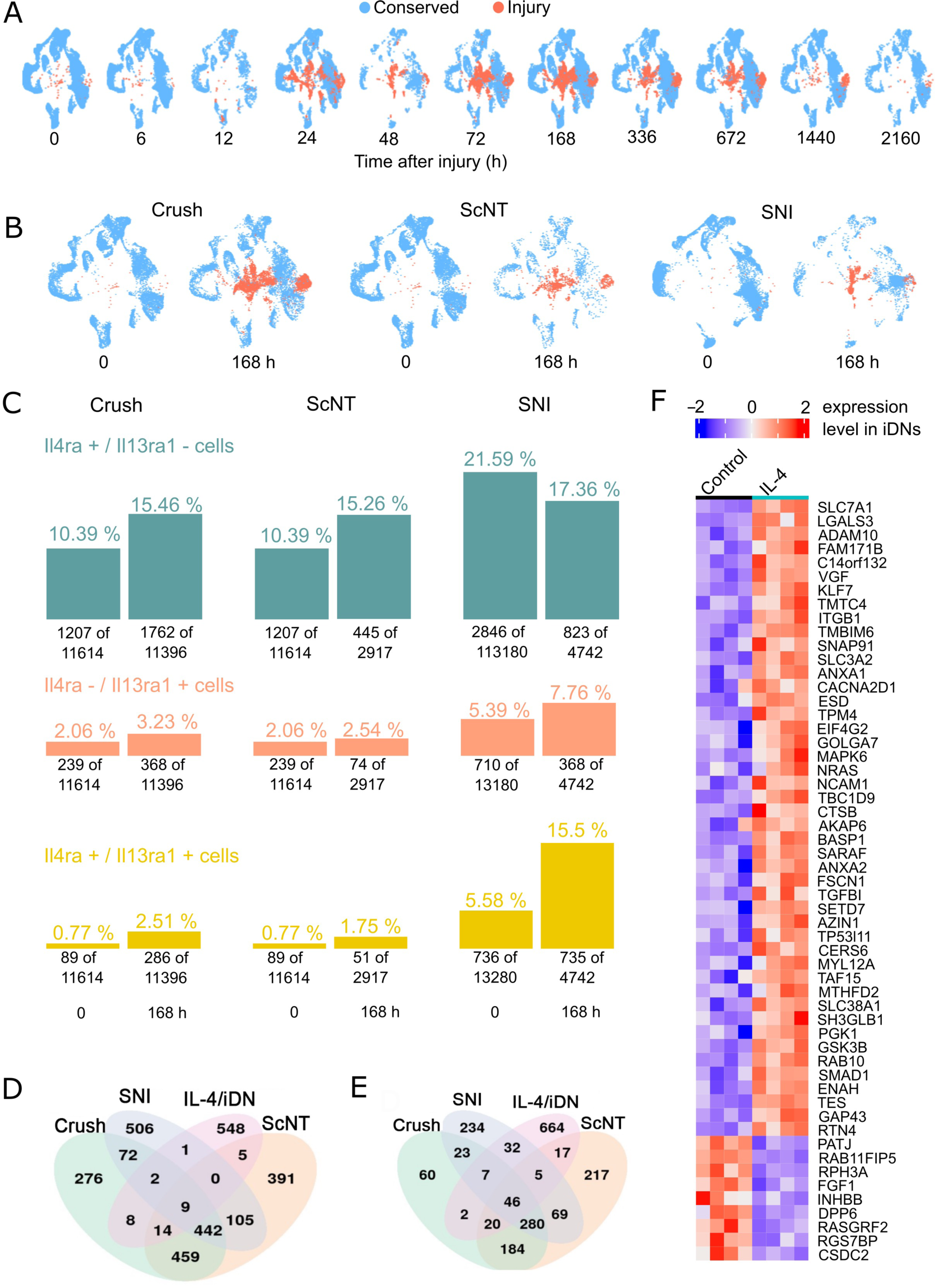
Il4ra and Il13ra1 expression in mouse injury models. (A) Publicly available mouse injury datasets with time courses after Crush injury, ScNT injury (GSE154659) and SNI (GSE155622) were integrated. Time point-specific changes in gene expression patterns reveal a common injury-cluster of cells not present under naïve conditions. (B) All three datasets provide data for the timepoint 168h (7d) post injury. (C) The absolute and relative numbers of Il4RA+/Il13RA1-, Il13RA1+/Il4RA- and Il4RA+/Il13RA1+ cells were compared between 0 and 168h post injury for all three injury datasets. (D, E) DEG analysis was performed between injury-induced neuronal- and the conserved clusters. The Venn plot shows the intersection of significantly up-and downregulated genes between all injury datasets and the Il-4 treated iDN DEGs. (F) The overall intersection revealed 46 commonly up and 9 commonly down-regulated genes illustrated for individual genes and the respective expression level in the iDN-IL-4 treated RNAseq dataset.

Relative gene expression analysis revealed significant (p<0.001) upregulation of Il4RA at 168h/7d post crush and SCNT as well as significant higher expression levels within the injury cluster compared to the conserved cluster. Il13RA1 was not found altered significantly. In the SNI data, no significant differences in l4ra expression levels were found between 0 and 168h nor conserved vs. injury cluster at day 7. However, significant (p<0.001) increased expression level of Il13RA1 was found in the injury cluster 168h/7d post injury (Supplementary Figure 6). To investigate the transcriptional profile of the emerging “injury “cluster and to potentially remodel it in-vitro, DGE-analysis was performed individually between cells from the “injury” cluster vs. “conserved” clusters. After FDR correction, significantly (p_adj_<0.05) regulated genes within the “injury” cluster were investigated: 622 upregulated and 1282 downregulated genes emerged 7d after crush injury, 838 upregulated and 1425 downregulated genes 7d after ScNT and 696 upregulated and 1137 downregulated genes 7d after SNI. Overall, 326 genes were commonly upregulated and 451 genes were downregulated in all three injury types (Fig. 4D-E). For the upregulated genes, GO enrichment analysis revealed the highest gene counts for axonogenesis (32 genes) and positive regulation of cell projection organization (26 genes) (Supplementary Figure 7).

In order to allow for comparison of mouse neurons and human iPSC derived iNocs, human DEGs were transformed into mouse-orthologues, and 793 orthologues were identified for the 932 human DEGs in iNocs. The comparison between IL-4 stimulated iNoc transcriptomic alterations and mouse scRNA data sets revealed an overlap of 75 upregulated and 33 downregulated genes, 88 upregulated and 28 downregulated genes between IL-4 iNocs and cells within a specific “injury” cluster in the sciatic nerve transection (ScNT) model as well as 90 upregulated and 12downregulated genes between IL-4-iNoc and a similar cluster in the spared nerve injury (SNI) model (Fig. 4D-E). GO analysis for the commonly upregulated genes (Fig. 4F) included genes involved in axonogenesis and axon guidance (*RTN4*, *GAP43*, *ENAH*, *RAB10*, *GSK3B*, *NCAM1*, *ITGB1*, *KLF7*), protein localization to cell periphery (*RAB10*, *MYL12A*, *GOLGA7*, *ITGB1*, *ADAM10*, *LGALS3*) and protein localization to plasma membrane (*RAB10*, *MYL12A*, *GOLGA7*, *ITGB1*, *LGALS3*). Genes that resembled ontologies like synapse organization (*GAP43*, *ADAM10*) or synapse assembly (*GAP43*) appeared already within previously mentioned ontologies like axonogenesis. These findings largely matched previous reports on presynaptic localization of IL4R (Hanuscheck et al., 2022), and suggested that IL-4 via Il4RA and possibly IL13RA1 through expressed downstream signalosomes controls the presynaptic components of iNoc neuronal networks. This could potentially be relevant for modelling neuroregenerative processes of peripheral nerves and their reconnections to spinal second order neurons.

## Discussion

In our study, IL-4 induced de novo formation of vGLUT1 immunoreactive boutons in developing iNocs which express all important components of the IL-4 receptor complex and signaling machinery at the iNoc stage. Inhibition of translational and cellular signaling components reduced this synaptogenic effect of IL-4. The involved distinct transcriptional changes in iNocs partially reflected injury-induced transcriptomic signatures retrieved in DRG neurons obtained from mouse models of peripheral nerve injury in vivo. The shared differentially expressed gene clusters were associated with axonogenesis and synapse formation contributing to regenerative processes both in peripheral nerve but possibly also for reconnecting primary afferent neurons to their projections in the spinal dorsal horn. Even a more general role of IL-4 in the control of presynaptic structures in developing and established neuronal networks may be anticipated.

The multifunctional cytokine IL-4 is mainly secreted by immune cells such as T-helper cells, eosinophils or mast cells (for recent review see: (Allen, 2023; Chakma & Good-Jacobson, 2023; Ding & Ge, 2022; Ji & Li, 2023) and binds to Il-4RA receptor homodimers or heterodimers consisting of IL-4RA and IL-13Ra1 subunits (Fujiwara, Hanissian, Tsytsykova, & Geha, 1997; Kammer et al., 1996; Van Dyken & Locksley, 2013). Both IL-4 and its receptors are expressed in the brain in neurons, oligodendroglia, astrocytes and microglia, and IL-4R expression increases after ischemic stroke indicative of a neuroprotective role (Amo-Aparicio et al., 2021; Bozinov et al., 2008; Fenn, Henry, Huang, Dugan, & Godbout, 2012; Lee, Koh, Lo, & Marchuk, 2018; T. F. Liu et al., 2003). About 25 % of neurons express IL-4RA, and an essential role of IL-4/IL-13 in the formation of inhibitory cortical synapses for normal social behavior is emerging for which a postsynaptic location of IL-4RA and IL-13RA1 is anticipated (Barron et al., 2023). In models of CNS injury, IL-4 protects neurons and induces recovery of injured neurons through neuronal IL-4RA (Walsh et al., 2015). In hippocampal neurons, IL-4 upregulates glutamatergic synaptic connections via presynaptic IL4R (Hanuscheck et al., 2022). We observed a similar presynaptic modulatory effect of IL-4 which increased numbers of vGLUT1 immunoreactive synaptic boutons in iNocs suggesting that IL-4 in general is a critically important regulator of presynaptic structures in the nervous system. In naive mouse DRG, we found that less than 25 % of neurons expressed IL4RA mRNA in all three datasets. After crush injury and ScNT, the percentage of neurons expressing Il4RA, Il13RA1 or both, Il4RA and Il13RA1 mRNA increased and this may be relevant for their ability to regenerate and reconnect to and reestablish glutamatergic synapses with their target neurons. The reported relative decrease of Il4ra only expressing cells 168h/7d after SNI (Fig. 4C) might indicate different repair-mechanisms that are specifically initiated in response to particular types of peripheral nerve injury. Although other cytokines such as IL-6 and TNF**a** are also involved in neuronal responses to injury promoting neuron survival and regeneration (Kummer, Zeidler, Kalpachidou, & Kress, 2021; Mi, 2008; Zeng, 2023) the effect on glutamatergic synapse formation was specific for IL-4 and moves this cytokine even more into focus as a promising novel tool to improve functional outcome after neuronal lesions.

In addition to glia and immune cell populations resident in the brain, IL-4RA protein is expressed in both excitatory and inhibitory neurons in cortex and hippocampus (Hanuscheck et al., 2022). Both brain regions are critically involved in cognitive performance and memory (Izquierdo et al., 2006; Jensen & Lisman, 2005). Previous studies report impairment of learning induced by a global IL-4 depletion which is rescued by IL-4–expressing T cells suggesting an indirect pathway putatively involving IL-4Ra expressing astrocytes (Derecki et al., 2010). The functional consequences of depleting IL-4RA specifically in neurons affect inhibitory as well as excitatory neuron populations and lead to controversial outcome if synaptic vesicle pools are reduced, postsynaptic currents altered, but excitatory drive increased in cortical networks of IL-4RA–deficient neurons (Hanuscheck et al., 2022). Improved exploratory behavior and locomotion, anxiety levels and mitigated fear learning through inhibitory synapses are reported in mice with a neuronal depletion of IL-4RA (GABAergic (Herz et al., 2021). Despite its effect on synaptic functions and although IL-4 causes neuron proliferation in hippocampus, a major role of IL-4/IL-4RA in neurodevelopment is not supported by studies in transgenic mice with a global depletion of IL-4RA which do not exhibit major deficits in cognitive performance and memory formation (Lee et al., 2018). However, with its dual importance for immune cells and neurons, IL-4 and the IL-4RA receptor are critically involved in the crosstalk between immune cells and neural progenitors governing plasticity and neurogenic capacity (Papadimitriou et al., 2018). Similar roles are emerging for the IL13Ra1 subunit of the heterodimeric receptor complex which as a synaptic protein modulates synapse physiology (Li et al., 2023). Activation of neuronal, synaptic IL-13Ra1 upregulates the phosphorylation of ionotropic glutamate receptors of both NMDA and AMPA subtype to increase synaptic activity (Li et al., 2023), and mice with a global depletion of IL-13 display similar deficits in learning behavior (Brombacher et al., 2017). In contrast to the well documented neuronal expression in the cns, the expression of IL4RA or IL13RA1 is not sufficiently well documented in primary sensory afferent neurons, but their presence is anticipated by functional studies reporting for example a nociceptor sensitizing effect of IL-13 (Barker et al., 2023). We found that naive primary afferents hardly expressed IL4RA or IL13RA1 mRNAs which was assigned to several neuron subtypes including peptidergic nociceptors. In contrast, increased IL4RA and IL13RA1 mRNA expression was revealed in an emerging “injury” cell cluster in three different nerve injury models with and without signatures of neuropathic pain supporting a critical role of IL-4 in the neuronal response to injury and the initiation of neuroregenerative processes (Umehara et al., 2020). After peripheral nerve injury in vivo, IL-4 expressing cells are recruited to the injury site, and IL-4RA expression increases in dorsal root ganglion neurons resulting in improved neurite and axon extension in vitro and in injured peripheral nerves in vivo (Pan et al., 2022). T cell–derived IL-4 protects and induces recovery of injured neurons by activation of neuronal IL-4 receptors potentiating neurotrophin signaling via AKT and MAPK pathways (Walsh et al., 2015) and primary neurons in culture respond to IL-4 with increased outgrowth (de Araujo-Martins et al., 2013; Gölz et al., 2006).

In iNocs, IL-4 mRNA was not retrieved, however, mRNAs coding for IL4RA and IL13RA1 receptors were expressed at relevant stages of iNoc differentiation and maturation together with STAT6 which is activated upon autophosphorylation of Jak1 and Jak2 (Murata, Obiri, & Puri, 1998), different isoforms of PKCs and translation factor EiF4E. IL-4 does not increase the total number of iNocs (Krakowiak et al., 2017; Nunan, Sivasathiaseelan, Khan, Zaben, & Gray, 2014) but induced de novo synthesis of vGLUT1 positive synaptic boutons indicative of a more mature neuronal phenotype. Our pharmacological approach provided evidence for the involvement of multiple signaling pathways that are activated by IL-4 in iNocs such as the mTOR pathway as published previously for bone marrow mesenchymal stem cells (Chen et al., 2015) or the contribution of protein kinases such as PKC or others (da Silva, Campello-Costa, Linden, & Sholl-Franco, 2008). Even translational processes may be involved in the signaling cascade as suggested by the inhibitory effect of EiF4E pharmacological Inhibition. The up-regulated gene clusters in iNocs stimulated with IL-4 partially overlapped with DEGs in nerve injury models indicative of regenerative processes and recovery. Our study promotes the importance of IL-4 and its receptors in the neuronal response to injury improving neuro-regeneration after nerve injury and the generation of glutamatergic synapses. With the current study we provide a model that not only can be used to increase our mechanistic understanding of pro-regenerative IL-4 effects in neurons but IL-4 can be added to culture media towards improving mature iNoc phenotypes in a human model system which can be useful as screening assay for future drug development. The emerging role of IL-4 acting via its receptors IL4RA and IL13RA1 is preserved in humans and offers novel opportunities not only as an interesting supplement to generate more mature phenotypes of iPSC derived neurons but also as a promising novel strategy for therapeutic interventions.

### Limitations of study

Increased development of glutamatergic synaptic boutons has been demonstrated in human iPSC derived neuronal cultures, however, not in the complexity of a live system such as spinal cord. Since for ethical reasons this is excluded to be verified in humans, verification currently only can be achieved in other species which are not fully compatible with humans or complex organoids consisting of iNocs synaptically connected to postsynaptic spinal neurons derived from iPSCs. These are currently still in their infancy but based on the discussed literature it is likely that confirmation can be obtained once they become available.

### Data Availability

The source code of all the bioinformatics can be accessed via github repository https://github.com/ZiDa20/il4_data_analysis. The RNAseq data set will be uploaded to GEO upon acceptance of the manuscript.

## Materials and methods

### Differentiation of iPSCs into nociceptors

iNocs were generated as previously described (Schoepf et al., 2020; Zeidler et al., 2021)In brief, we used the human iPSC line SFC840-03-01 (STBCi026-B) generated within the StemBANCC project (part of Innovative Medicines Initiative) based on a commercially available fibroblast line (Lonza) from a healthy 67-years old female donor (Morrison et al., 2015)(Morrison et al. 2015). The cells were routinely cultured on cell culture plastic ware (Greiner) coated with a murine ECM preparation (Matrigel™ at 8 µg/cm2; Corning). Cells were fed three times a week with mTeSR1 plus medium (STEMCELL Technologies) and passaged twice a week by dissociation in 0.02% EDTA. The cells were used from the earliest available passage number over 20 passages maximum. For differentiation into iNOCs, iPSC cultures were singularized in Accutase® (Gibco) and seeded onto Matrigel® at 70.000 cells/cm2 in the presence of 10µM Rho-kinase inhibitor Y-27632 (MedChemExpress). Differentiation was initiated by dual SMAD inhibition (100nM LDN-193198, 10µM SB431542; Selleckchem), leading to neural crest like cells followed by an overlapping inhibition of GSK3β, VEGF and Notch signaling (3i inhibition: 3µM CHIR-99021, 10µM SU5402, 10µM DAPT; SelleckChem), and finally long-term maintenance in Neurobasal® medium (Gibco) supplemented with N2 (1:50; Gibco), B-27 (1:100; Gibco) and the neurotrophic factors GDNF (25ng/ml), **D**β-NGF (25ng/ml), and BDNF (10ng/ml; all from Preprotech).

### Cytokine treatment and Immunofluorescence microscopy

Cryopreserved d12 iNocs were plated in N2/B-27 Neurobasal medium including growth factors (see previous paragraph) at a density of 1,7 x 10^4^ cells on Matrigel-coated 13mm glass coverslips in 24-well plates. At d15 cells were treated with IL-4 (20 ng/ml, LOT130-094-117, Miltenyi Biotec), IL-6 (100 ng/ml) or TNF-**α** (10 ng/ml) for 3h and 24h before fixation with 4% paraformaldehyde in PBS at room temperature for 10 minutes. Untreated controls were included for both time points. Fixed iNocs were treated with 5% normal goat serum in PBS supplemented with 0.2% BSA and 0.2% Triton X-100 for 30 minutes. Primary antibodies were applied at 4°C overnight in a humidified chamber and detected by fluorochrome-conjugated secondary antibodies (Alexa goat anti-rabbit A594 (#A32740), goat anti-mouse A488 (#A32723), 1:4000; Invitrogen). Primary antibodies used were anti-vGLUT1 (1:2000, rabbit polyclonal, Synaptic Systems (SySy, A#135302) (Schöpf et al., 2021)), anti-Tuj1 (1:600, mouse monoclonal, R&D Systems, #MAB1195, (Quarta et al., 2014)). Nuclei were counterstained with DAPI (4’,6-diamidino-2-phenylindol) 1:10.000, (Thermo Fisher Scientific). Images were recorded using an Axioimager 2 Microscope (Carl Zeiss Microscopy) with cooled CCD camera (SPOT Imaging Solutions). Average fluorescence intensities were quantified using Metaview ® software (Molecular Devices, LLC) with the line scan plug-in to quantify fluorescent intensity along a defined line cutting the individual synaptic structures.

For inhibitor treatments D12 iNocs were seeded and cultured as described in the previous paragraph. On d15, cells were pretreated for 1h with STAT 6 inhibitor AS1517499 (1µM; Sigma Aldrich SML1906), eIF4E/eIF4G interaction inhibitor 4EGI-1 (40 µM; Merck 324517), mammalian Target of Rapamycin (200 nMmTOR) inhibitor Rapamycin (200nM; Merck 553210) and PKC inhibitor Bisindolylmaleimide (1µM; Merck 203290), followed by 24h of IL-4 (20 ng) stimulation. All inhibitors were dissolved in DMSO, and DMSO concentration (0,05%) was kept constant across all conditions. Cells were fixed with 4% PFA and further used for IF experiments. Inhibitor treated conditions were compared to DMSO only controls or IL-4 treated iNocs.

To quantify synaptic boutons, triple stained IF images (TUJ1/vGLUT1/DAPI) were processed in Metaview®. Images were gray value thresholded. Area Limit Filters were set between 0 and 60 to include individual vGLUT1+ clusters whereas nuclei were excluded from the measurements. Object size was further controlled by using the measure calibrate distance function adjusted to the corresponding objective: Following a final object control in the histogram, object data was assessed and IMA counts for individual vGLUT+ clusters measured. In order to distinguish between vGLUT1+ clusters transported along axons and vGLUT1+synaptic boutons only TUJ1/vGLUT1 double positive boutons fulfilling the threshold 0.5 µm-5 µm in a defined area of a 40x image were assessed by the manually count object mask of Metaview®. Proper synaptic structure was further evaluated by line scan analysis for the vGLUT1/594 and TUJ1/488 fluorescent signal. Thus, fluorescent amplitudes were measured along a line of 3 µm cutting the individual presynaptic bouton. As a control for the stable number of plated iNoc neurons DAPI staining was assessed between different treatment conditions. Average fluorescent intensities were background subtracted, plotted in MS excel and finally illustrated with Graph Pad Prism and Adobe Photoshop 2023.

### RNA extraction and sequencing

RNA extraction was performed from eight biological samples containing each 500k cells (four replicates untreated (ut) controls, four replicates of 24h IL-4 (20 ng/ml) treated neurons) using Quantseq kit followed by high-throughput Illumina sequencing. Samples (frozen pellets of iPSC derived neurons) were shipped on dry ice, stored at −80°C until processing and total RNA was extracted using the Lexogen SPLIT RNA Extraction Kit (User Guide Version 008UG005V0320) according to the manufacturer’s guidelines. Samples were characterized by UV-Vis spectrophotometry (Nanodrop2000c, Thermo Fisher), RNA integrity was assessed on a Fragment Analyzer System using the DNF-471 RNA Kit (15 nt) (Agilent). Sequencing-ready libraries were produced using QuantSeq 3’ mRNA-Seq V2 Library Prep Kit with UDI following standard procedures, as outlined in the respective User Guide (191UG444V0111): Indexed library preparation was performed to allow for multiplexed sequencing. For library preparation, 400 ng of total RNA samples were used as an input. Prepared libraries were quality controlled on a Fragment Analyzer device (Agilent), using the HSDNA assay. Concentration of obtained libraries was quantified using a Qubit dsDNA HS assay (Thermo Fisher). A sequencing-ready pool of indexed libraries was prepared according to these quantifications. Sequencing was performed on an Illumina platform.

### RNA-seq data sets and bioinformatics analysis

Raw sequencing reads quality and phred quality scores were assessed manually using *fastqc* (v. 0.11.9). The overall raw read length was 100 bp. The raw sequencing reads were deduplicated to reduce the impact of amplification bias using *umi-tools* (vers. 1.1.4). Low quality bases and adaptor sequences were trimmed using *bbduk* (BBMap v. 38.18) with trimming parameters set to kmer size = 13, use of shorter kmers enabled and the minimum kmers threshold was set to 5. Quality trimming was enabled to remove reads with a read length smaller than 20 nucleotides and base pairs with a quality score < 10. Deduplicated sequencing reads were aligned against the human reference genome (Gr38) using STAR-aligner (v. 2.7.0). Remaining PCR duplicates were removed in a second deduplication step performed using umi-tools. The raw transcript counts per gene were processed into a count matrix using feature counts of the Rsubread package (v 2.12.3) in R (v. 4.2.2). Low expressed genes with a threshold <10 counts were removed from the count matrix and differentially regulated genes were identified using DESeq2 (v. 1.38.3) (Love, Huber, & Anders, 2014). To correct for multiple testing, false detection rate (FDR) correction was performed using the Benjamini-’Hochberg algorithm (Love et al., 2014) resulting in adjusted p-values (p-adj). Gene-Ontology enrichment analysis (GO) was performed using ClusterProfiler (v 4.6.2) for genes with p-adj<0.0 1. Before GO, genes were ranked according to the product of log2foldchange*(-log10(p-adj)).

Human (GSE168243) and mouse (GSE155622, GSE154659) DRG sequencing data were downloaded from GEOData and preprocessed identical using Seurat v.4.3.0 and R 4.2.2. Preprocessing comprised batch-wise log normalization (NormalizeData) and scaling (ScaleData) of the raw counts. Appropriate integrations anchors were selected for all batches and integration was performed using canonical correlation analysis based on the integration anchors. On the integrated object, principal component analysis (PCA), clustering and neighbor identification was performed. Cluster annotation for peptidergic neurons (PEP1, PEP2), neurofilament positive clusters (Ab-LTMR, Ad-LTMR and Proprioceptors), a non-peptidergic-cluster, pruriceptors, c-LTMRs, putative c-LTMRs and injury was performed according to marker genes as introduced (Renthal et al., 2020).

For each dataset, differential gene expression (DGE, FindAllMarkers()) was performed between the injury cluster and all other cells at selected time points. Benjamini-Hochberg FDR corrected significantly differentially expressed genes (p_adj < 0.05) were joined for downstream analysis. Venn-diagrams were created using library Venn Diagramm 1.7.3 and gene ontology enrichment analysis was performed using Cluster Profiler library 4.6.2. Genes given into GO analysis were ranked according their adjusted p-value and the average log2 fold change expression avg_log2FC*(-log(p-adj) value.

### Microfluorimetric Ca^2+^ measurements

Ca^2+^ measurements of iNocs were performed as we previously described (Camprubí-Robles et al., 2013). Cells were loaded with 3 µM of Fura-2 AM (Invitrogen) in extracellular solution (ECS) containing: 145 mM NaCl, 5 mM KCl, 2 mM CaCl2, 1 mM MgCl2, 10 mM D-glucose and 10 mM HEPES with pH adjusted to 7.3 with NaOH, for 30 min at 37°C in 5% CO2. Subsequently, cells were washed with ECS. Calcium imaging was performed on an IX71 microscope (Olympus) with 20x/0.85 N.A. oil-immersion objective. The Omicron LEDHUB system (Omicron-Laserage Laserprodukte GmbH, Rodgau-Dudenhofen, Germany) was used for excitation of Fura-2 at 340 nm and 380 nm wavelengths and emission was detected at 510 nm. F340/F380 excitation ratio was calculated after background correction. Baseline was recorded over 10 s prior to each treatment. To elucidate treatment induced Ca^2+^ transients in an unbiased way an increase of 10% over baseline F340/F380 ratio was applied as threshold. The percentage of active cells was calculated and compared for each treatment and time-point.

## Contributor Roles

D.Z., C.L.S. and M.K: Conceptualization

D.Z., C.L.S., G.K., M.Z. and T.K.: Methodology; Validation; Formal analysis; Investigation

D.Z. and M.Z.: SoftwareResources; Data curation

D.Z., C.L.S., G.K. and M.K.; Writing – original draft preparation; Writing – review & editing

D.Z., C.L.S.: Visualisation

M.K.: Supervision; Project administration; Funding acquisition

## Impact statement

Convergent evidence from morphological and transcriptomic analyses supports a more general role of IL-4 in the formation of glutamatergic synapses in differentiating neurons derived from human induced pluripotent stem cells.

## Acknowledgements

We are grateful for funding of this work by the Austrian Research Fund FWF (Project I5087) to MK.

---

**Supplementary Figure 1 IL-4 specific role in synaptic bouton formation in iDNs.** (A-D) iDN neurons were treated with different cytokines (IL-6, TNF**α**) in comparison to IL-4 treatment for 24h. (A-B) IL-4 (20ng/ml) robustly upregulated the number of TUJ1+ boutons in comparison to untreated controls. (C-D) 24h treatment with IL-6 (100ng/ml) and TNF**α** (10ng/ml) did not cause a significant change in the number of TUJ1+ boutons; (E) Quantification of TUJ1+ boutons by IMA measurements; (F) The number of iDN somata did not change upon the different treatments;

**Supplementary Figure 2** iNocs functionally respond to glutamate. Ca^2+^ transients evoked by chemical nociceptor stimuli (A) Time-series analysis of substance induced Ca^2+^ transients following administration of extracellular solution (ECS/BASELINE), 100 µM glutamate, and 25 mM KCL (dark grey) was quantified and visualized. (B) Relative response magnitude for each positive cell was determined and substance induced reaction was compared qualitatively. (C) The stacked bar charts indicate the percentage of positively reacting and non-reacting cells upon administration of glutamate and KCL; (D) Genes for putatively involved glutamate receptors were queried in NOCICEPTRA and illustrated in a heatmap.

**Supplementary Figure 3** Cells per area did not change between individual treatments (inhibitors: STAT6i (1µM), eIF4Ei (40µM), RAP (200 nM), BIM (1 µM)) compared to ut-control and IL-4 treated iDNs.

**Supplementary Figure 4** Representative images of iDNs (four samples control, four samples IL-4 treated iDNs analyzed by bulk RNA sequencing.

**Supplementary Figure 5** (A) Comparison of relative proportion of cells per neuronal subtype at different timepoints post injury within the integrated RNAseq data. (B) *Il4RA* expression levels within neuronal subtypes before and 168h post injury compared between Crush, ScNT and SNI data.

**Supplementary Figure 6** Gene expression levels of *Il4RA* and *Il13RA1* were compared between all cells from naïve and 168h/7d after injury as well as between conserved and injury labeled cells at 168h/7d after injury.

**Supplementary Figure 7** Gene Ontology enrichment analysis for the intersecting genes significantly upregulated (FDR correct p-value < 0.05) in the injury cluster at 168h post injury of the respective injury type.

