## Supplementary figures and images for "IL-4 shaping glutamatergic synapse like structures for more mature iPSC derived neuron phenotypes"

### supplementary figures 1-7

Suppl. Figure 1

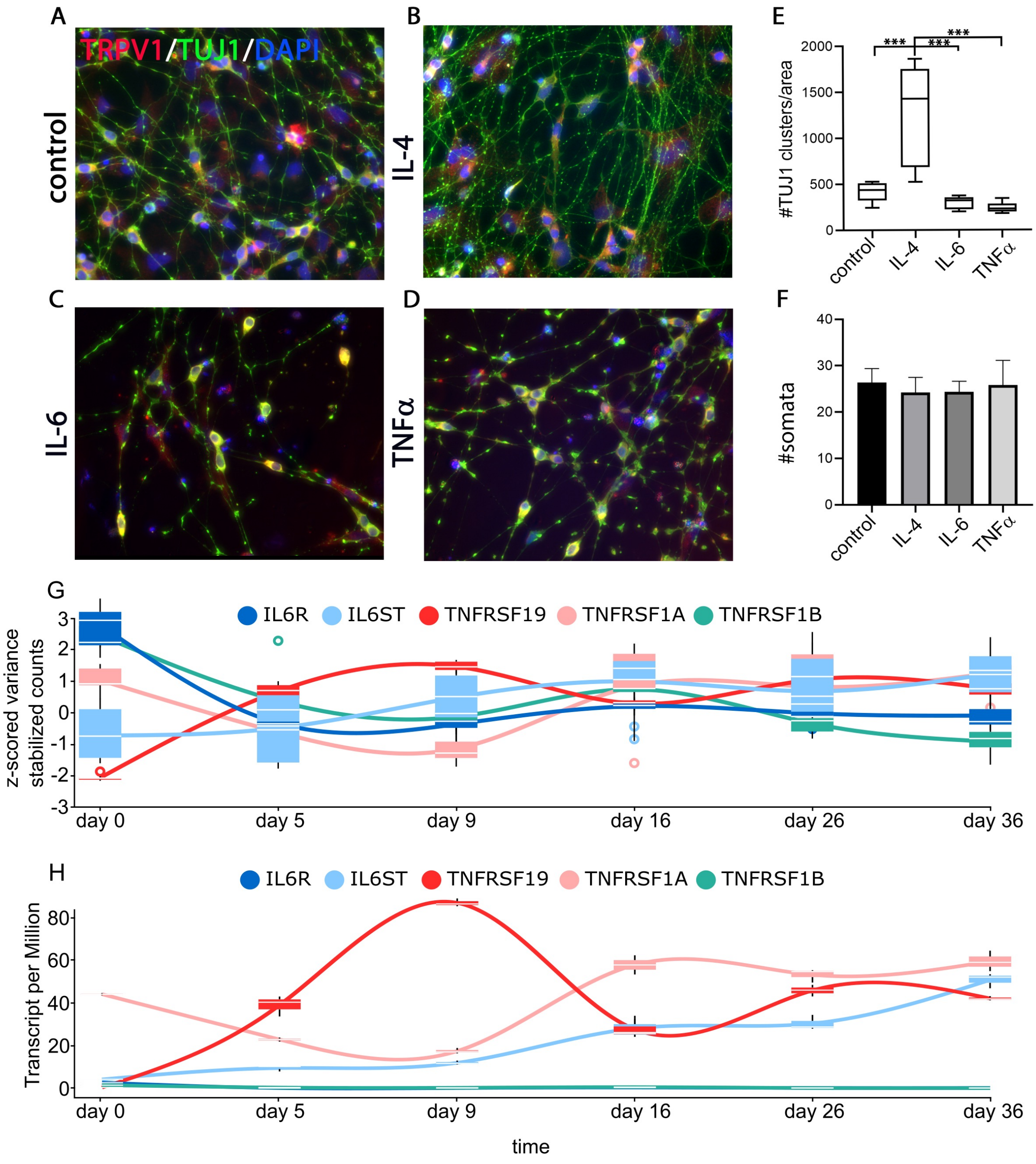

Suppl. Figure 2

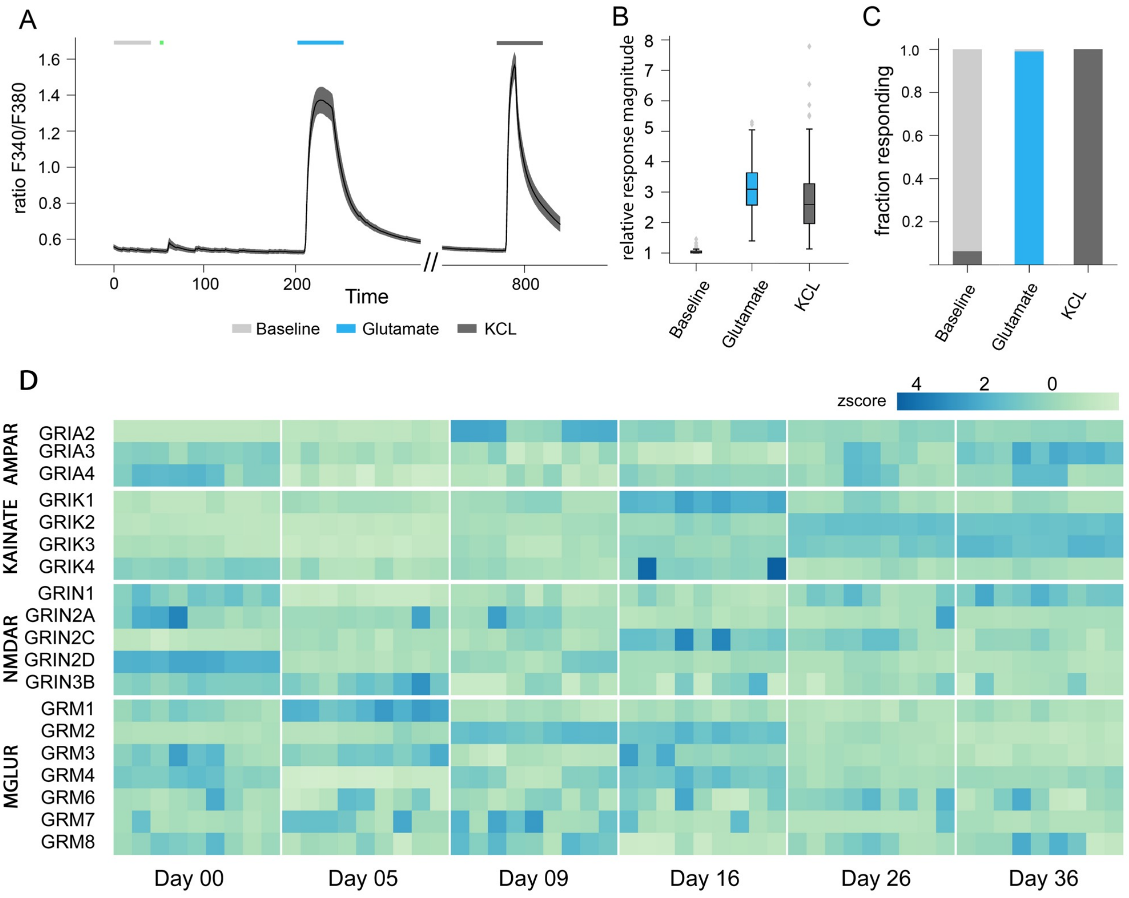

Suppl. Figure 3

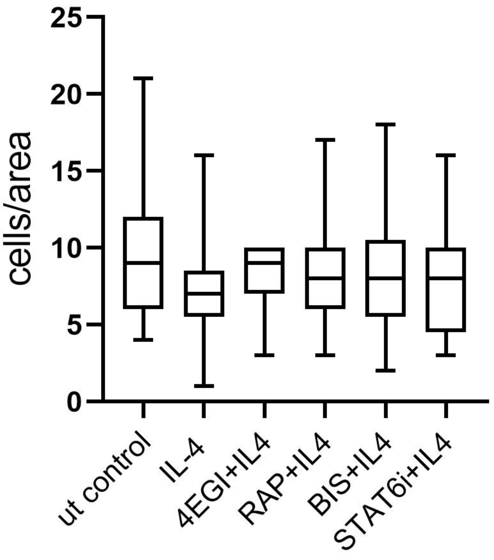

Suppl. Figure 4

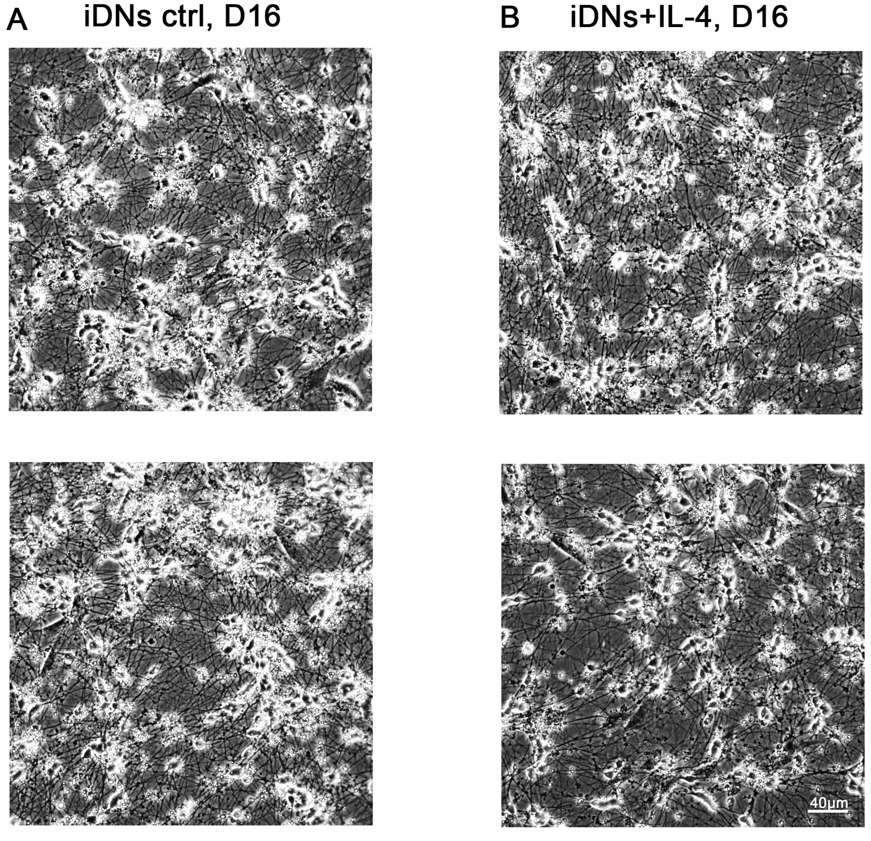

Suppl. Figure 5

A

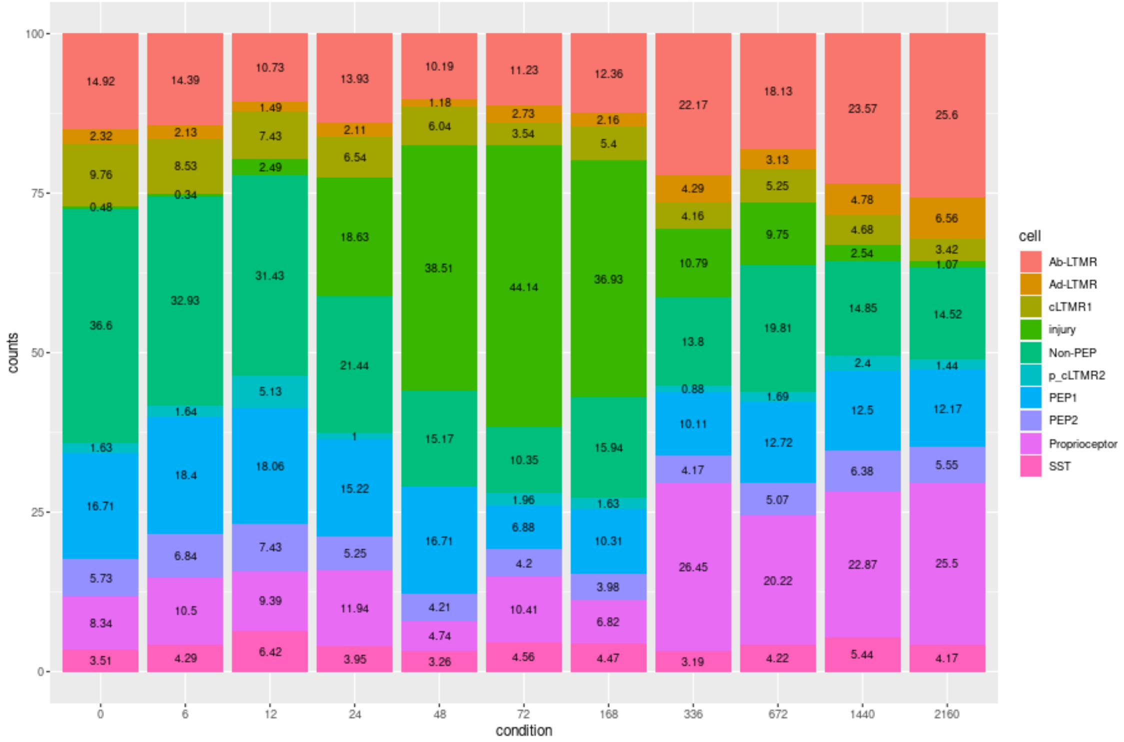


B

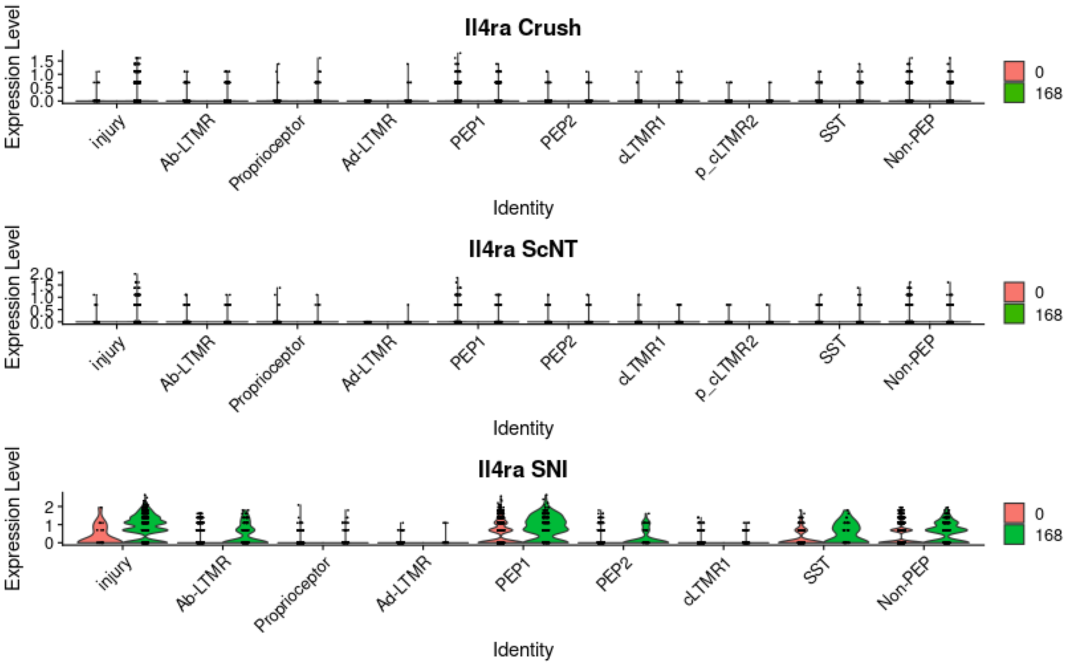

Suppl. Figure 6

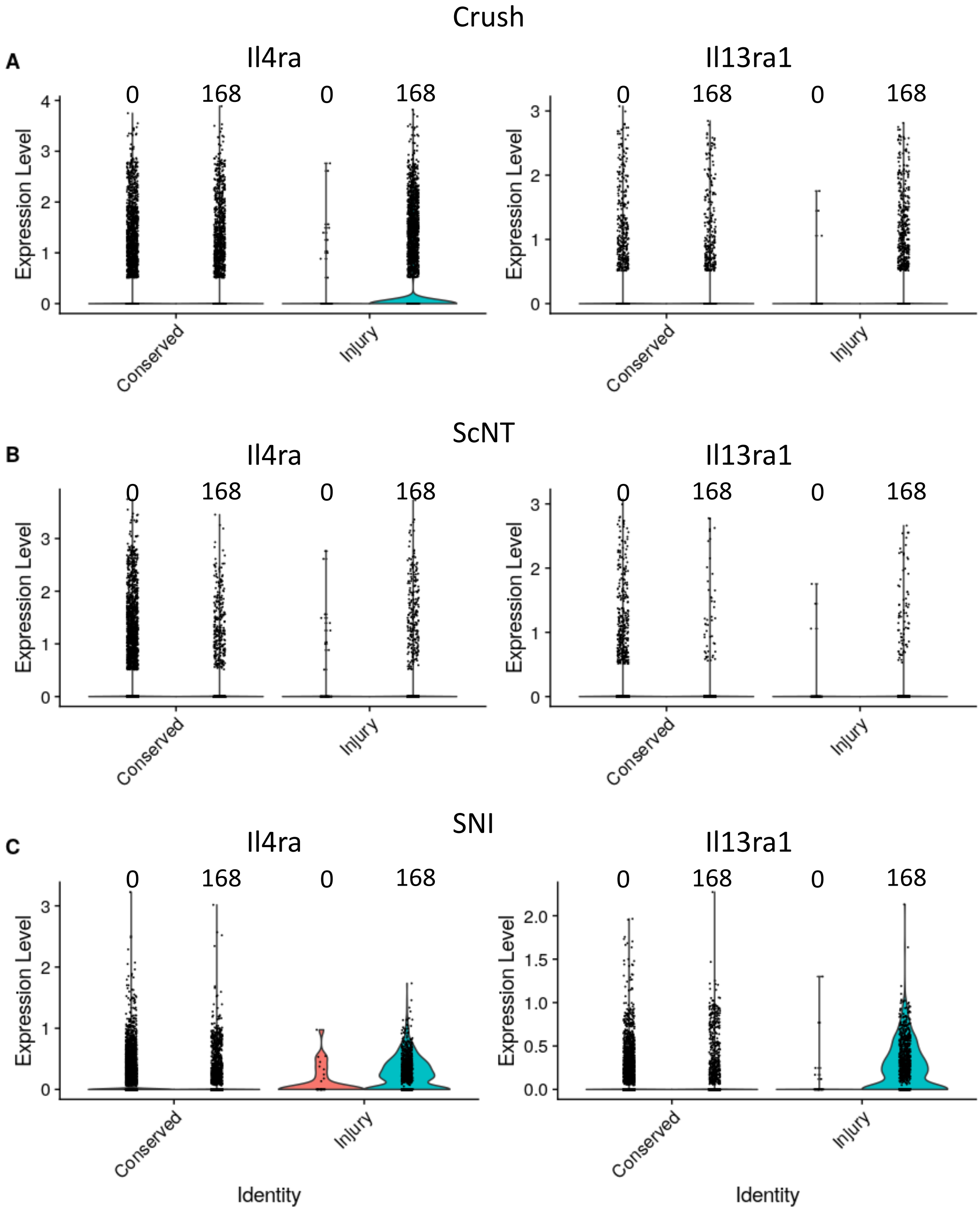

Suppl. Figure 7

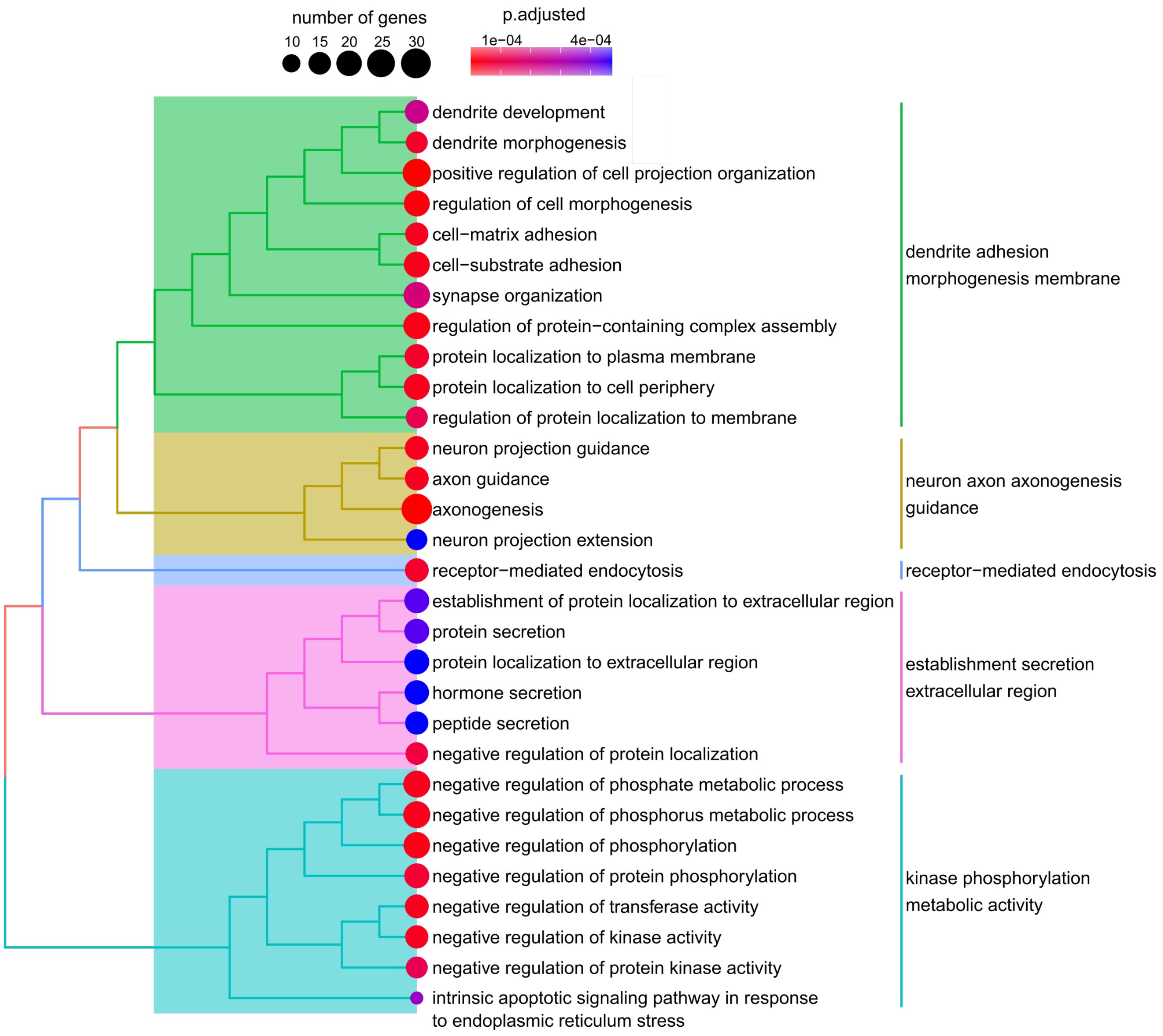
